# Multi-locus characterization and phylogenetic inference of *Leishmania spp.* in snakes from Northwest China

**DOI:** 10.1101/511162

**Authors:** Han Chen, Jiao Li, Junrong Zhang, Jinlei He, Jianhui Zhang, Zhiwan Zheng, Dali Chen, Jianping Chen

## Abstract

**Background:** Leishmaniasis caused by protozoan parasite *Leishmania* is a neglected disease which is endemic in the northwest of China. Reptiles were considered to be the potential reservoir hosts for mammalian Leishmaniasis, and *Leishmania* had been detected in lizards from the epidemic area in the northwest of China. To date, few studies are focused on the natural infection of snakes with *Leishmania*.

**Methods:** In this study, 15 snakes captured from 10 endemic foci in the northwest of China were detected *Leishmania spp.* on the base of mitochondrial *cytochrome b*, *heat shock protein 70* gene and *ribosomal internal transcribed spacer 1* regions, and identified with phylogenetic and network analyses.

**Result:** In total, *Leishmania* gene was found in 7 snakes. The phylogenetic inference trees and network analysis suggests that the species identification was confirmed as *Leishmania donovani*, *Leishmania turanica* and *Leishmania sp*.

**Conclusion:** Our work is the first time to investigate the natural *Leishmania spp.* infection of snakes in the northwest of China. Mammalian *Leishmania* was discovered in snakes and the reptilian *Leishmania* was closely related to the clinical strains both prompt the importance of snakes in the disease cycle. To indicate the epidemiological involvement of snakes, a wide sample size in epidemic area and the pathogenic features of reptilian *Leishmania* promastigotes are recommended in the future research.

## Introduction

Leishmaniasis is a neglected disease caused by infection with flagellate protozoan parasite *Leishmania* of the family *Trypanosomatidae*. There are three main forms of leishmaniasis: visceral or kala-azar (VL), cutaneous (CL) and mucocutaneous (MCL). Over 20 *Leishmania* species known to be infective to humans are transmitted by the bite of infected sand flies of which at least 98 species were proven or probable vectors worldwide [1]. An estimated 2 million new cases and 50 000 deaths occur in over 98 countries annually [2]. In China, which is one of 14 high-burden countries of VL [3], there exists three epidemiological types: anthroponotic VL (AVL), mountain-type zoonotic VL (MT-ZVL), and desert-type ZVL (DT-ZVL) [4]. Acknowledging the information on the epidemiology including the vector and animal reservoir hosts of the disease is better to understand the disease and its control.

In view of the topography in the northwest of China, DT-ZVL is found largely in the oases and deserts including the southern and eastern of Xinjiang Uygur Autonomous Region, the western part of the Inner Mongolia Autonomous Region, and northern Gansu province [5]. Surveillance of the disease reflects that DT-ZVL is persistence with sporadic outbreaks and still not under control now [5]. Sand flies (*Phlebotominae*) were recognized as the transmission vector in the traditional sense, while ticks (*Ixodoidea*) had been reported to be the vector in the transmission of canine VL [6–9]. In addition to humans, there are various kinds of hosts, mainly mammals such as canids, hyraxes and rodents, which could be the source of infection as the reservoirs. Because transmission occurs in a complex biological system involving the human host, parasite, vectors and in some cases an animal reservoir host, the control of Leishmaniasis is pretty complex. As for the desert in the northwest of China, there is no consistent agreement regarding dissemination of the actors playing the key roles in leishmaniasis.

Moreover, reptiles, mainly lizards, were found harboring *Leishmania* parasites with controversies in the part of spreading the disease [10]. Blood cells of lizards containing amastigotes were first found by Chatton and Blanc in *Tarentola mauritanica* from southern Tunisia [11] and then several cases from the same lizard species were reported in the next decades [12]. Wenyon provisionally named the leptomonad flagellates in gecko as *Leishmania tarentolae* [13]. Killick-Kendrick recognized *Sauroleishmania* as a separate genus for the leishmanial parasites of reptiles [14], later DNA sequence-based phylogenies had clearly placed it within the *Leishmania* genus as a secondarily derived development from the mammalian species [15–17]. Most studies were focused on the infection of *Leishmania* parasites in lizards, but few were in snakes except Belova and Bogdanov cultured promastigotes from the blood of five species of snakes in the Turkmen S.S.R. [14,18].

In reality, the detection of reptilian *Leishmania* is rare in China. Zhang et al. was the first team to detect *Leishmania* via molecular methods from the lizards captured in the northwest of China [19]. Interestingly, besides reptilian *Leishmania*, mammalian *Leishmania* species (*L. donovani* and *L. tropica*) were involved despite *Sauroleishmania* was considered not to infect humans or other mammals [20,21]. This may support the potential reservoir role of lizards for human leishmaniasis.

Similarly, we employed the highly sensitive polymerase chain reaction (PCR) to detect the presence of *Leishmania spp.* in snakes. The sequences of various genetic markers have been used successfully to infer the phylogenetic relationships within *Leishmania* including the sequences of DNA polymerase α [15], RNA polymerase II [15], 7SL RNA [22–24], ribosomal internal transcribed spacer [25,26], mitochondrial *cytochrome b* gene [27,28], *heat shock protein 70* gene [29,30] and the *N-acetylglucosamine-1-phosphate transferase* gene [31]. Therefore, this study was based on three genetic loci, i.e., *cytochrome b* (*cyt b*), *heat shock protein 70* (*Hsp70*), and *internal transcribed spacer 1* (*ITS1*) region to genetically characterize *Leishmania spp.* detected in 15 snakes, and conduct tree-based species delimitation to infer the phylogenetic positions by comparison with some representative *Leishmania* sequences retrieved from GenBank. It was the first time to survey the infected snakes of *Leishmania* in the northwest of China and try to reveal the primary evidence on the epidemic role of snakes for human leishmaniasis.

## Materials and methods

### Study area and sampling

15 snakes identified as 3 species by morphological characters were captured alive by hand and snake hooks at 10 sites across the endemic foci of VL in the northwest of China (Table 1). These sites are located in arid desert areas with the altitude ranging from 537 to 1222 m above sea level. Franchini [32,33] seems to have observed rare amastigotes from the liver of lizards, and Shortt & Swaminath [34] cultured a strain *Leishmania* on NNN (Novy-MacNeal-Nicolle) medium from the liver of lizards. Consequently, a part of the liver was taken from the abdominal cavity of each snake to detect the presence of *Leishmania* via PCR. In total, 15 liver specimens were collected in Eppendorf tube and stored at −20°.

**Table 1.**
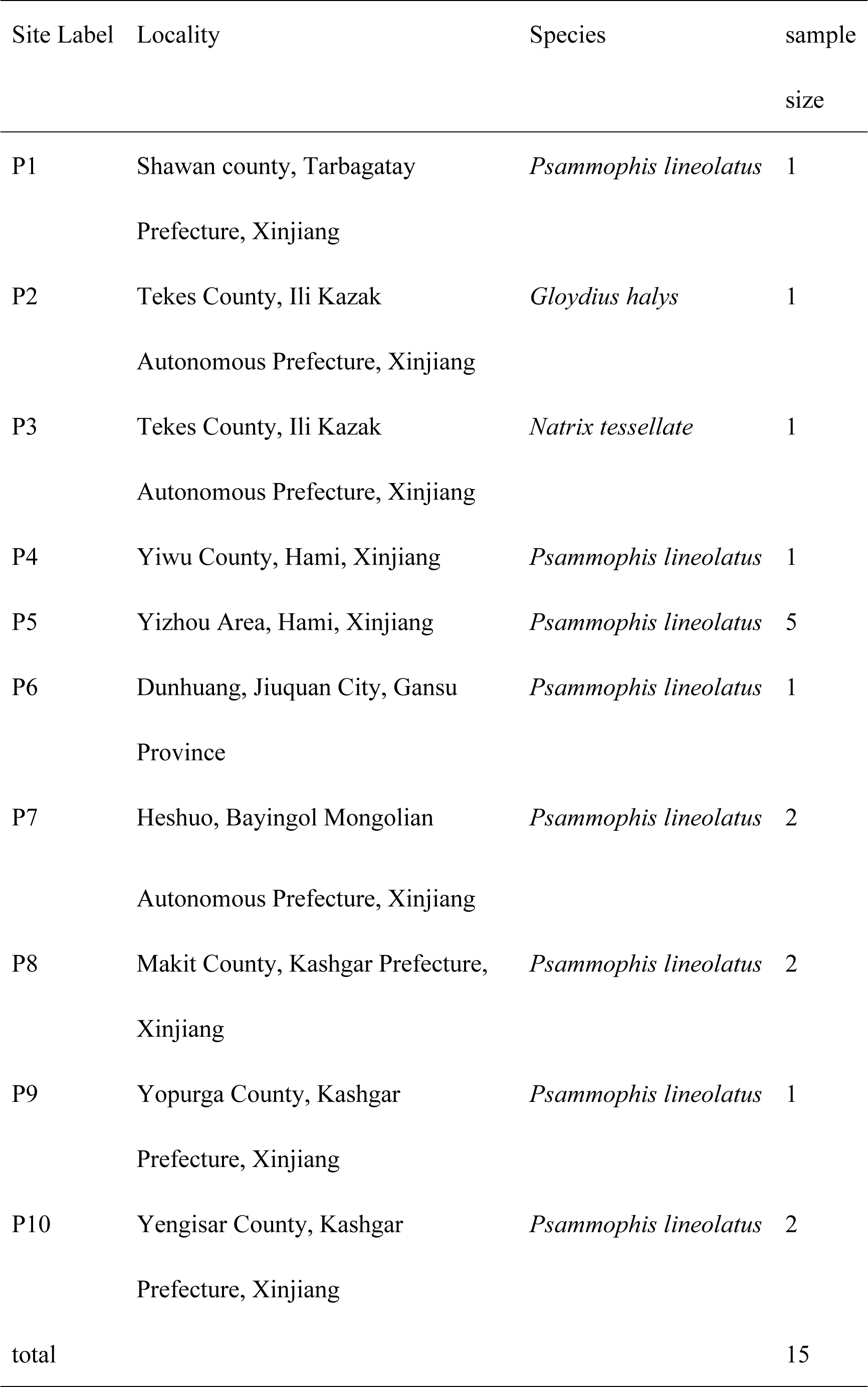
List of sampling localities, snake species and sample size in this study.

All surgery was performed under sodium pentobarbital anesthesia, and all efforts were made to minimize suffering. The protocol was approved by the ethics committee of Sichuan University (Protocol Number: K2018056).

### DNA extraction, amplification, cloning and sequencing protocols

Total genomic DNA was extracted from liver tissues using the commercial kit, TIANamp^®^ Genomic DNA Kit (TIANGEN Bio, Beijing, China) following the protocols of the manufacturer. PCR primers specific for the *Leishmania* synthesized by Tsingke Biological Technology Co., Ltd (Chengdu, China) were used to amplify *cyt b* [35], *Hsp70* [36] and *ITS1* [37] gene fragments by PrimeSTAR Max DNA polymerase (TaKaRa Bio, Shiga, Japan) according to the manufacturer’s instruction. We picked *Leishmania* strain MHOM/CN/90/SC10H2 as the positive control while preparing the mixture, and the negative control was treated without template DNA. The PCR products were purified by excision of the band from agarose gel using the Universal DNA Purification Kit (TIANGEN Bio, Beijing, China), and ligated into a pGEM^®^-T vector (Promega Co., Madison, USA) with added the poly (A) tails. After transformation of *Escherichia coli* DH5α competent cells with the ligation product and X-gal blue-white screening, 8 positive colonies were selected by PCR using the plasmid primers T7 and SP6, and grown in LB/ampicillin medium. DNA was extracted and sequenced at Tsingke Biological Technology Co., Ltd (Chengdu, China). The chromatograms were validated and assembled in BioEdit v7.2.6 [38].

### Sequence alignment and phylogenetic analyses

GenBank searches were performed to initially identify the species of the original DNA sequences using BLASTn (https://blast.ncbi.nlm.nih.gov/Blast.cgi). And all the nucleotide sequences generated in this study have been submitted to the GenBank database. The sequences were multiple-aligned with a set of *Leishmania* strains of each locus examined in this study retrieved from the GenBank using ClustalW of MEGA (Molecular Evolutionary Genetic Analysis v7.0.26 [39]) with its default option and refined manually. The alignments were further trimmed to exclude regions with missing data and then distinct haplotypes were defined by DAMBE v7.0.1 [40,41]. The evolutionary history was inferred by phylogenetic tree construction using Bayesian inference (BI). Gaps were treated as missing data and each haplotype was treated as a taxon in the analyses. The program PartitionFinder v2.1.1 [42] was used to select the most appropriated substitution model of all phylogenetic analyses. Bayesian analyses were carried out using the program MrBayes v3.2 [43] which the trees were started randomly. 2 parallel sets of 4 simultaneous Monte Carlo Markov chains (3 hot and 1 cold) were run for 10,000,000 generations until convergence was reached (stopval = 0.01) and the trees were sampled for every 1000 generations. Temperature heating parameter was set to 0.2 (temp = 0.2) and burn-in was set to 25% (burninfrac = 0.25). The results of Bayesian analyses were visualized in FigTree v1.4.3 (available at http://tree.bio.ed.ac.uk/software/figtree/).

### Network reconstructions

Due to the limit of bifurcating tree to represent intraspecific gene evolution, haplotype networks may more effectively portray the relationships among haplotypes within species. The phylogenetic networks were constructed by using the Median Joining (MJ) network reconstruction method in NETWORK v5.0.0.3 (available at http://www.fluxus-engineering.com/sharenet.htm) [44,45].

## Results

### *Leishmania* infection of the captured snakes in the study area

Through preliminary determination of identity using BLASTn, the total prevalence of *Leishmania* DNA detected via PCR was 46.7% (7/15). 5 (33.3%) snakes were detected as positive for *cyt b*, 4 (26.7%) snakes for *Hsp70* and 2 (13.3%) for *ITS1*. Considering the sensitivity of different loci, two samples from P1 and P2 were amplified successfully in all three loci, while other samples were detected by only one locus in this study. The snakes from P1, 2, 3, 4, 5, 6, 9 were *Leishmania*-positive of all the 10 foci as shown in Fig 1.

**Figure 1.**
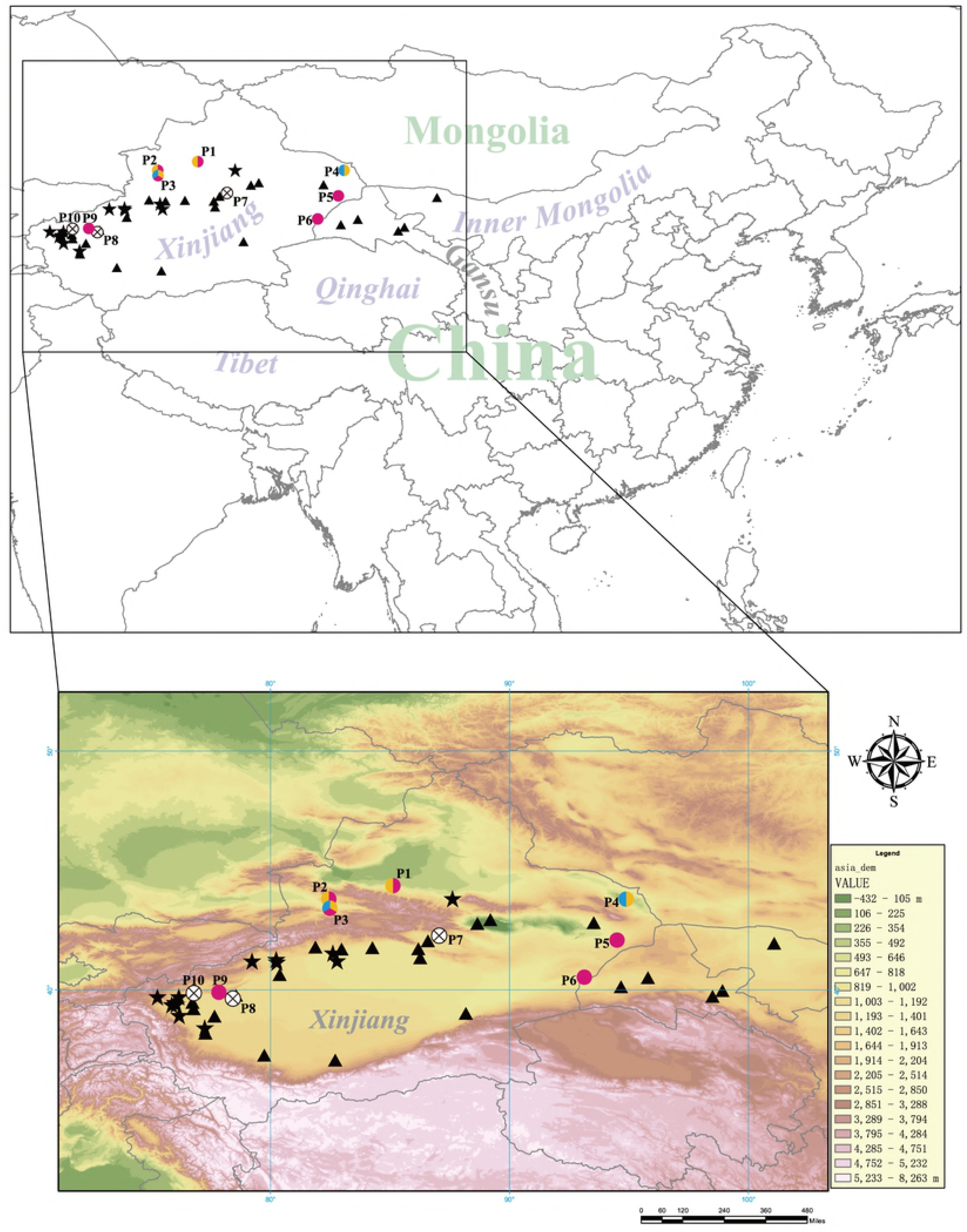
Sampling sites of snakes in Northwest China, along with the current foci of endemicity of anthroponotic visceral leishmaniasis (AVL) and desert-type zoonotic visceral leishmaniasis (DT-ZVL) in China. The site numbers P1-P10 correspond to those in Table 1. Solid five-pointed star (★) represents the endemic focus of AVL and solid triangle (▲) represents the endemic focus of DT-ZVL [4,5]. The species of *Leishmania* detected from the samples are shown for different colors (yellow: *L. donovani complex*, blue: *L. turanica*, pink: *Leishmania sp.*)

### Sequence characteristics

After the haplotypes were defined, 10 *cyt b* sequences, 7 *Hsp70* sequences and 10 *ITS1* sequences were obtained in this study and deposited in the GenBank database under Accession No. MK330198-MK330207, MK330208-MK330214 and MK300708-MK300717, respectively (Table 2). PCR amplification of each target locus resulted in amplicons of the expected sizes as follows: *cyt b* (543 bp), *Hsp70* (738-767 bp), and *ITS1* (328-334 bp). The GC contents were 23.4% for *cyt b*, 64.9% for *Hsp70* and 41.8% for *ITS1*, respectively. Thus, both their lengths and their GC contents are within the range of corresponding sequences of other *Leishmania* species.

**Table 2.**
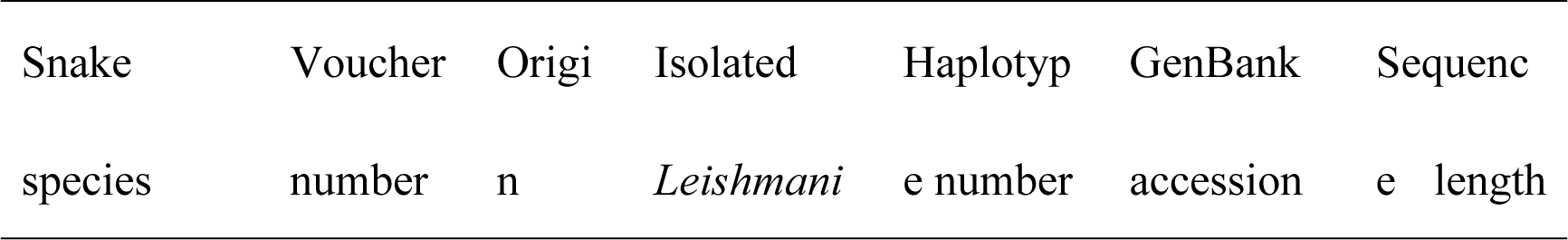
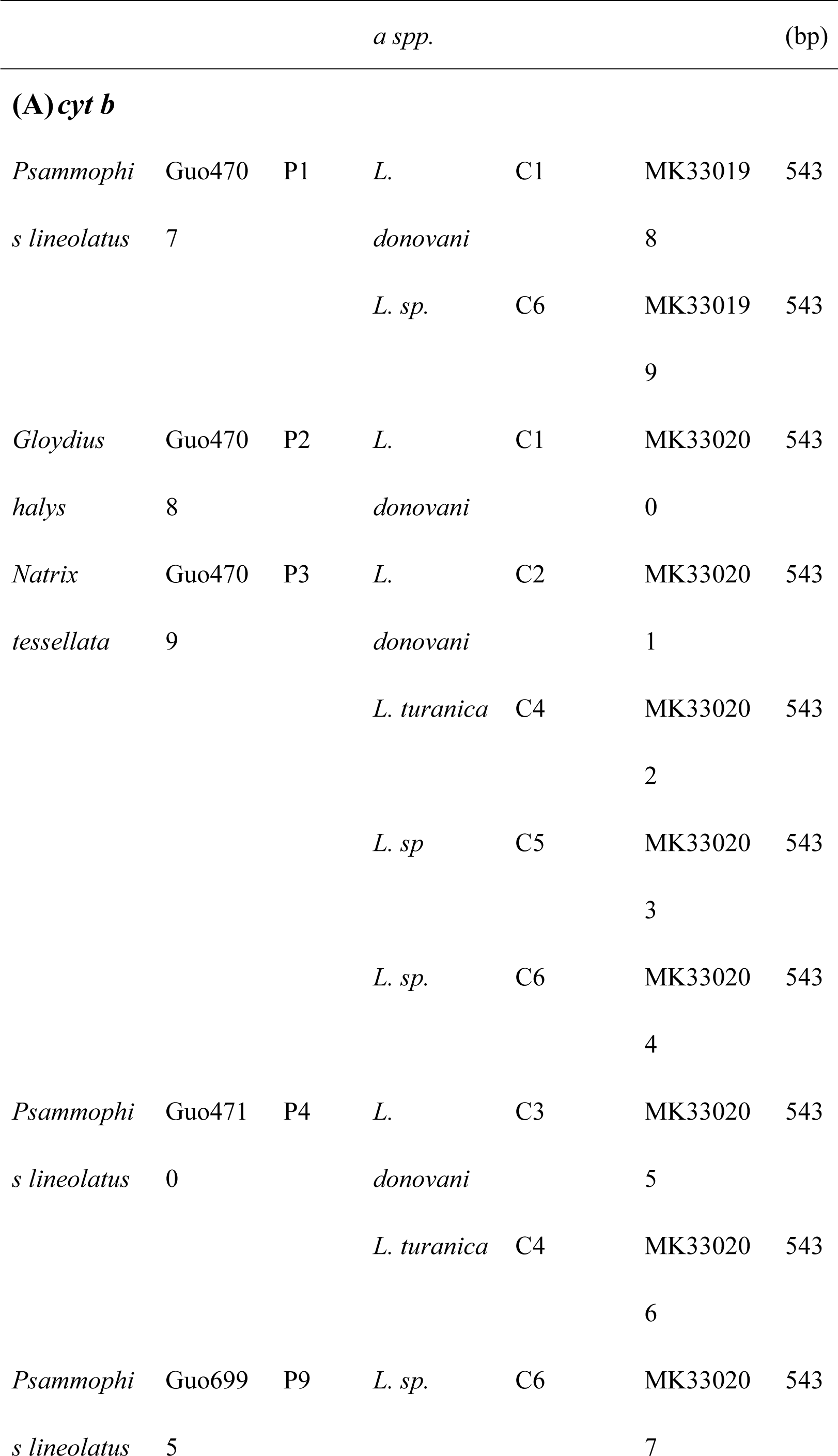
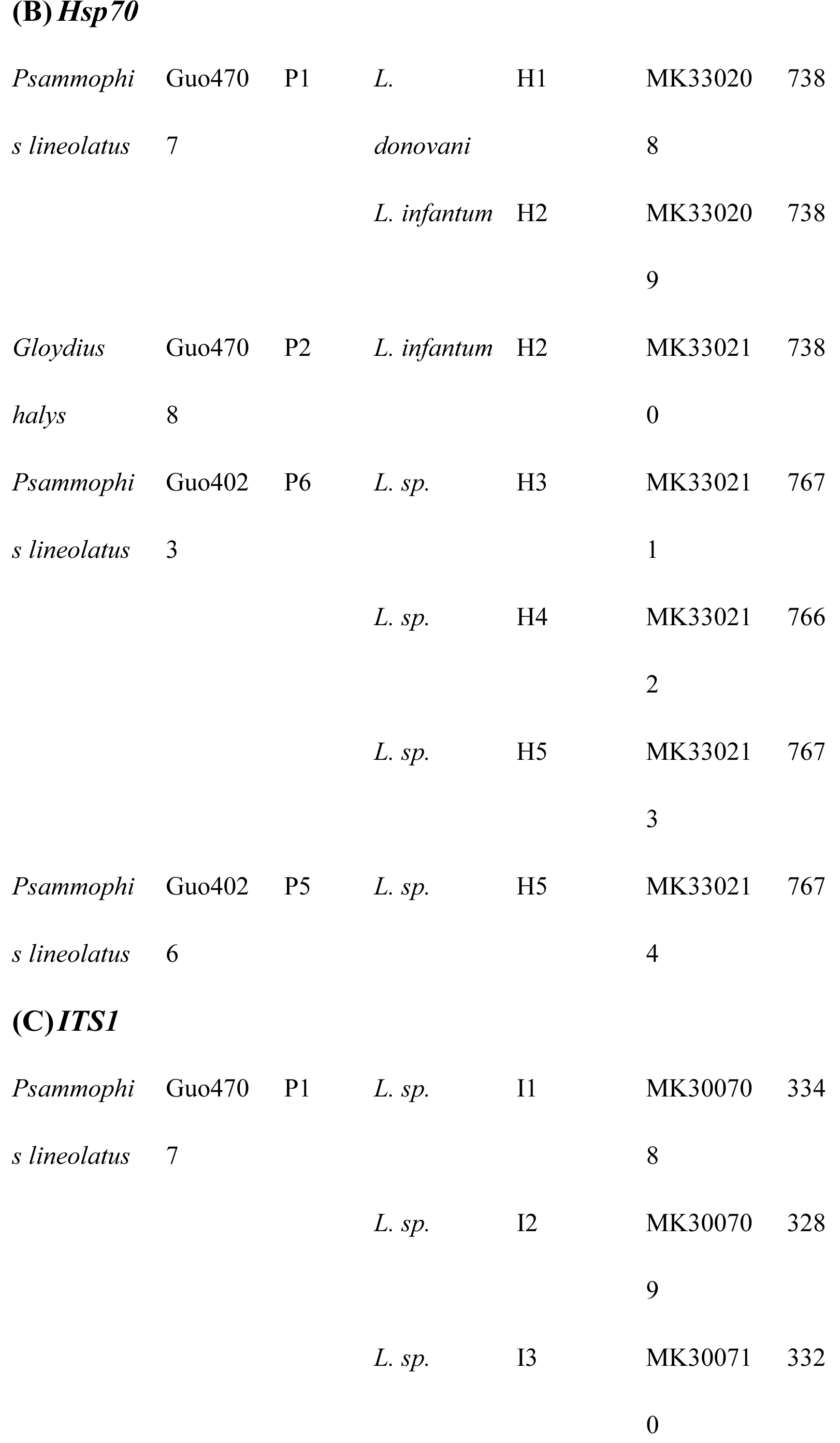
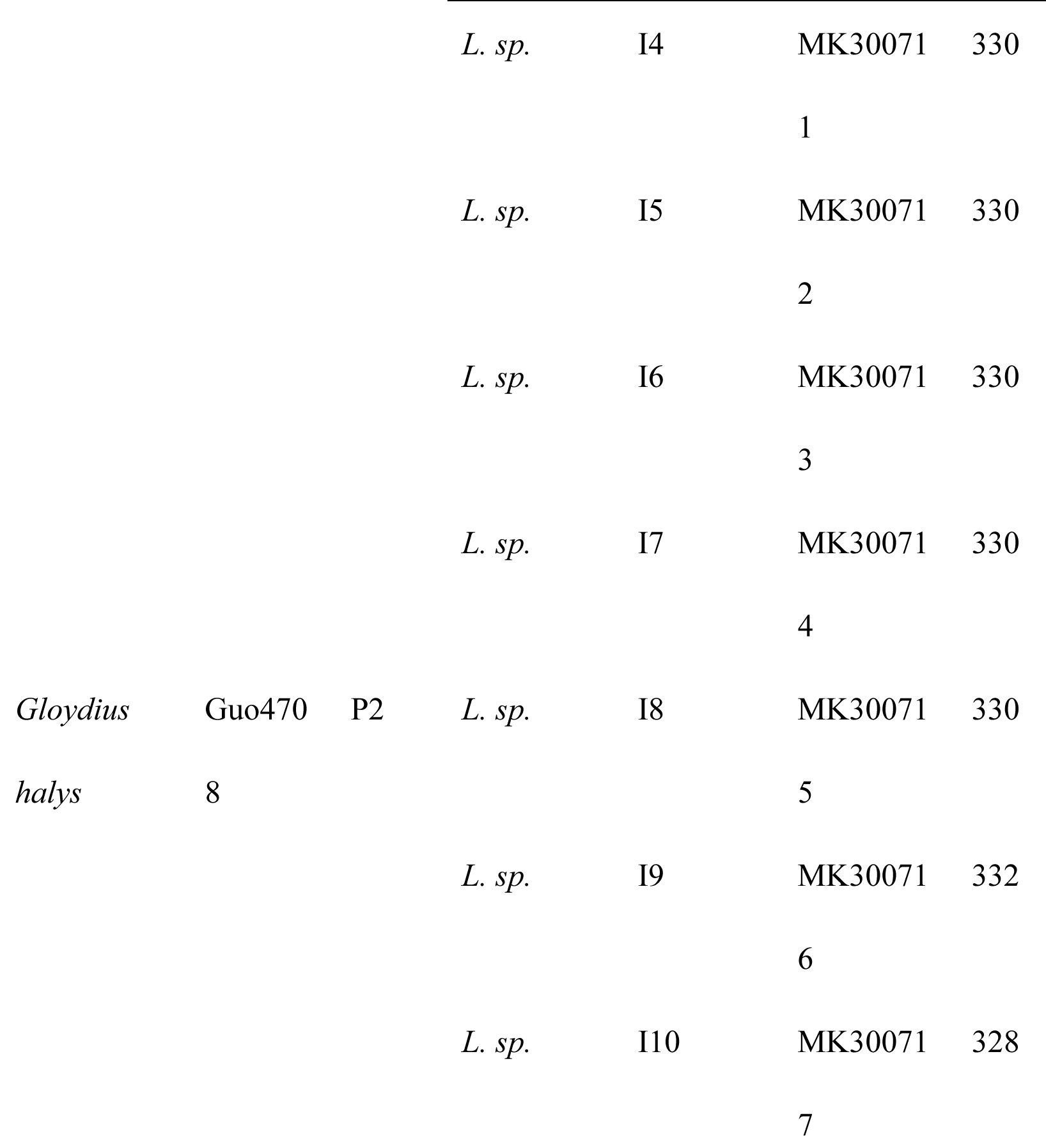
List of the lizard samples, origin, isolated *Leishmania spp.* and GenBank accession numbers.

| Snake species | Voucher number | Origin | Isolated *Leishmania* | Haplotype number | GenBank accession | Sequence length |
| --- | --- | --- | --- | --- | --- | --- |

| *a spp.* |  |  |  |  |  | (bp) |
| --- | --- | --- | --- | --- | --- | --- |
| (A)*cyt b* |  |  |  |  |  |  |
| *Psammophi* | Guo470 | P1 | *L.* | C1 | MK33019 | 543 |
| *s lineolatus* | 7 |  | *donovani* |  | 8 |  |
|  |  |  | *L. sp.* | C6 | MK33019 | 543 |
|  |  |  |  |  | 9 |  |
| *Gloydus* | Guo470 | P2 | *L.* | C1 | MK33020 | 543 |
| *halys* | 8 |  | *donovani* |  | 0 |  |
| *Natrix* | Guo470 | P3 | *L.* | C2 | MK33020 | 543 |
| *tessellata* | 9 |  | *donovani* |  | 1 |  |
|  |  |  | *L. turanica* | C4 | MK33020 | 543 |
|  |  |  |  |  | 2 |  |
|  |  |  | *L. sp* | C5 | MK33020 | 543 |
|  |  |  |  |  | 3 |  |
|  |  |  | *L. sp.* | C6 | MK33020 | 543 |
|  |  |  |  |  | 4 |  |
| *Psammophi* | Guo471 | P4 | *L.* | C3 | MK33020 | 543 |
| *s lineolatus* | 0 |  | *donovani* |  | 5 |  |
|  |  |  | *L. turanica* | C4 | MK33020 | 543 |
|  |  |  |  |  | 6 |  |
| *Psammophi* | Guo699 | P9 | *L. sp.* | C6 | MK33020 | 543 |
| *s lineolatus* | 5 |  |  |  | 7 |  |
| **(B) *Hsp70*** |  |  |  |  |  |  |
| *Psammophi* | Guo470 | P1 | *L.* | H1 | MK33020 | 738 |
| *s lineolatus* | 7 |  | *donovani* |  | 8 |  |
|  |  |  | *L. infantum* | H2 | MK33020 | 738 |
|  |  |  |  |  | 9 |  |
| *Gloydus* | Guo470 | P2 | *L. infantum* | H2 | MK33021 | 738 |
| *halys* | 8 |  |  |  | 0 |  |
| *Psammophi* | Guo402 | P6 | *L. sp.* | H3 | MK33021 | 767 |
| *s lineolatus* | 3 |  |  |  | 1 |  |
|  |  |  | *L. sp.* | H4 | MK33021 | 766 |
|  |  |  |  |  | 2 |  |
|  |  |  | *L. sp.* | H5 | MK33021 | 767 |
|  |  |  |  |  | 3 |  |
| *Psammophi* | Guo402 | P5 | *L. sp.* | H5 | MK33021 | 767 |
| *s lineolatus* | 6 |  |  |  | 4 |  |
| **(C) *ITS1*** |  |  |  |  |  |  |
| *Psammophi* | Guo470 | P1 | *L. sp.* | I1 | MK30070 | 334 |
| *s lineolatus* | 7 |  |  |  | 8 |  |
|  |  |  | *L. sp.* | I2 | MK30070 | 328 |
|  |  |  |  |  | 9 |  |
|  |  |  | *L. sp.* | I3 | MK30071 | 332 |
|  |  |  |  |  | 0 |  |
|  |  |  | *L. sp.* | I4 | MK30071 | 330 |
|  |  |  |  |  | 1 |  |
|  |  |  | *L. sp.* | I5 | MK30071 | 330 |
|  |  |  |  |  | 2 |  |
|  |  |  | *L. sp.* | I6 | MK30071 | 330 |
|  |  |  |  |  | 3 |  |
|  |  |  | *L. sp.* | I7 | MK30071 | 330 |
|  |  |  |  |  | 4 |  |
| *Gloydus* | Guo470 | P2 | *L. sp.* | I8 | MK30071 | 330 |
| *halys* | 8 |  |  |  | 5 |  |
|  |  |  | *L. sp.* | I9 | MK30071 | 332 |
|  |  |  |  |  | 6 |  |
|  |  |  | *L. sp.* | I10 | MK30071 | 328 |
|  |  |  |  |  | 7 |  |

### Phylogenetic relationship

Prior to the phylogenetic analyses, the most adequate models of nucleotide substitution were selected by PartitionFinder v2.1.1: GTR +G and GTR+I+G for *cyt b*, HKY+I for *Hsp70*, K80+G for *ITS1*. Using Bayesian inference method, the trees showed similar phylogenetic topology for all three loci supported by posterior probability (PP) values.

Regarding the BI tree inferred from each locus, the phylogenetic analyses inferred from the aligned matrix of *cyt b* (Fig 2 (A)) separated two distance clades, the *Viannia* isolates apparently formed an independent clade (PP=1.0) outside the clusters of those other species (PP=1.0), while the subgenera *Leishmania* and *Sauroleishmania* were still separated into different branches (PP=1.0 and 0.56, respectively). The isolates C1, C2 and C3 were clustered with *L. donovani* and *L. infantum* (PP=1.0). The isolate C4 was shared an identical *cyt b* sequence to a *L. turanica* isolate and they were grouped with *L.arabica*, *L. major*, *L. tropica*, *L. aethiopica* and *L. killicki* in a subclade (PP=0.91) separated with *L. donovani complex*. The isolate C6 shared identical sequences with Chinese *Leishmania sp.* isolates from Yang et al. [26] was clustered in a separated clade (PP=0.96) together with C5 and other *Leishmania sp*.

**Figure 2.**
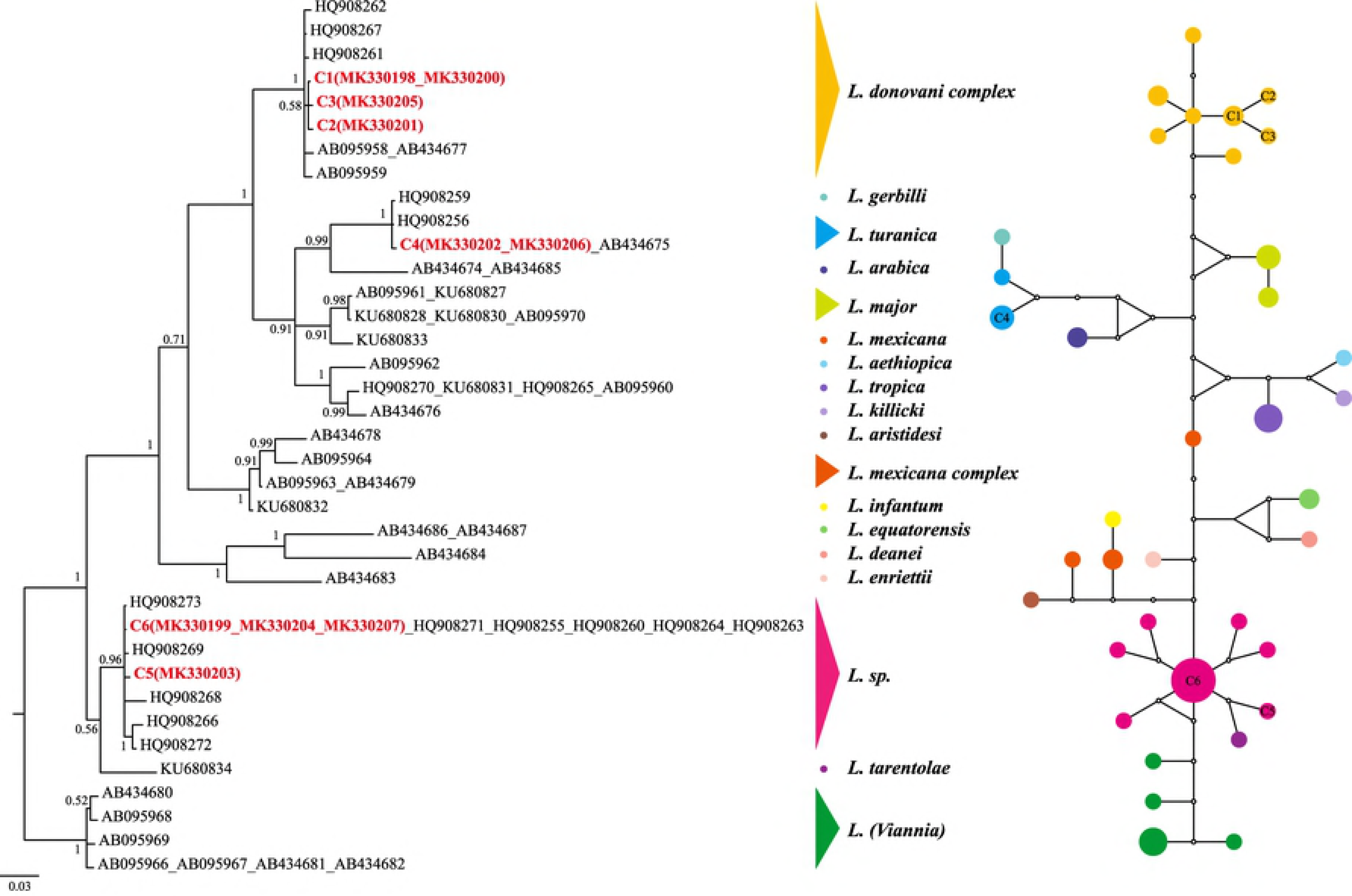
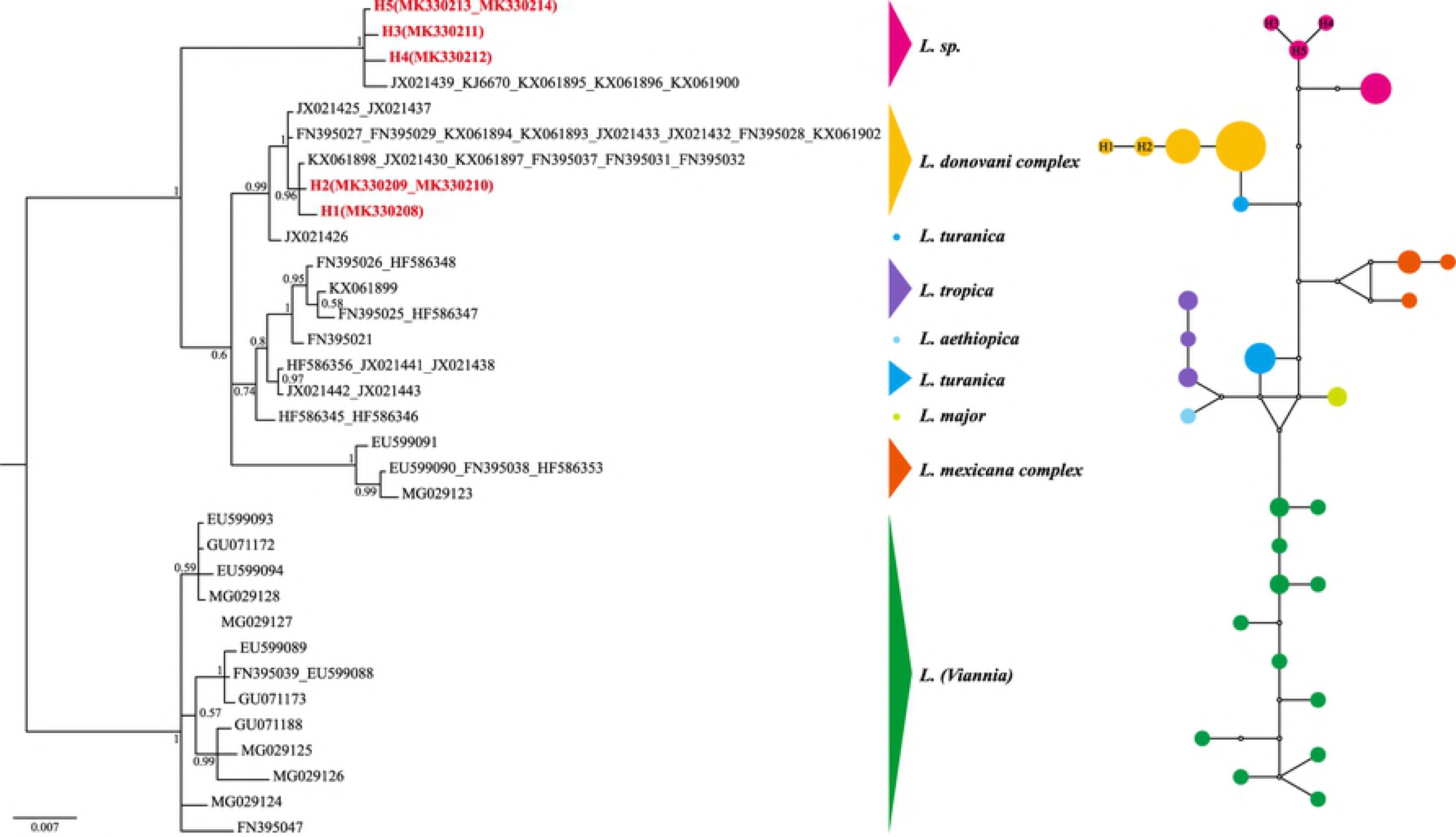
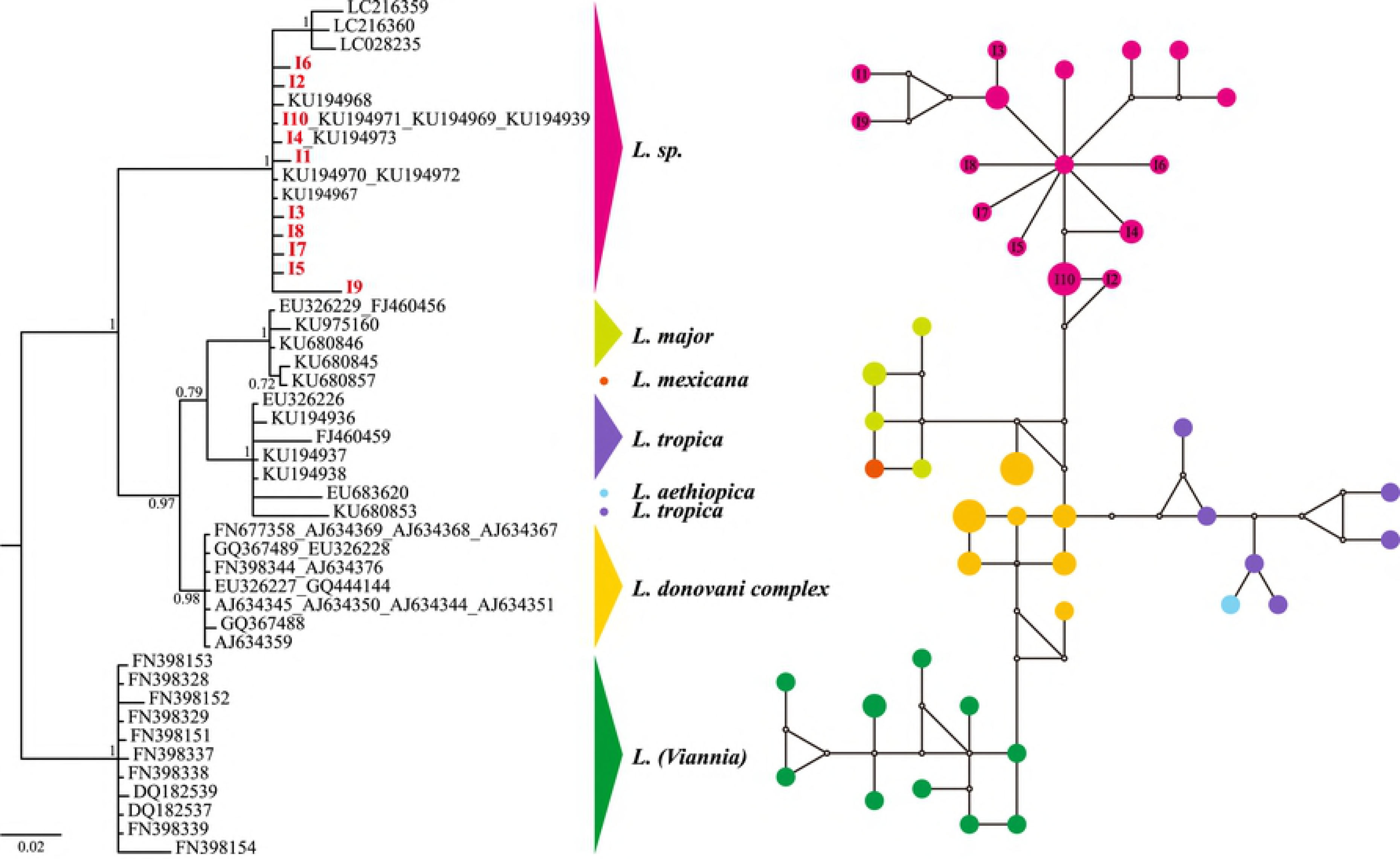
The majority-rule consensus tree (midpoint rooted) of three genetic loci inferred from Bayesian inference by using MrBayes v.3.2, and the corresponding median-joining networks were implemented by NETWORK v5.0.0.3: (A) *Cyt b*, (B) *Hsp70*, (C)*ITS1*.

The *Hsp70* analysis (Fig 2 (B)) showed a similar structure to the phylogenetic tree of *cyt b*: *Viannia* (PP=1.0), *Sauroleishmania* (PP=1.0) and *Leishmania* (PP=0.6). The isolates H1 and H2 searching by BLASTn (Megablast) revealed over 98% identity with

*L. donovani* and *L. infantum* respectively, while both of them were nested within *L. donovani complex*. In this subclade, the isolates of other species of *Leishmania* subgenus (except *L. mexicana* clustered by the PP 1.0) were not in reciprocally monophyletic groups. The isolates H3, H4 and H5 were clustered in a separated clade (PP=1.0) with *Leishmania sp.* from China.

For the *ITS1*-5.8S region, the phylogenetic tree as Fig 2 (C) shown was constructed by three clades with high support (PP=1.0, 0.97 and 1.0, respectively), corresponding to the three subgenera *Viannia*, *Leishmania*, and *Sauroleishmania*. All the 10 fragments from snakes formed a strongly supported cluster with *Leishmania sp.* isolated from lizards [19] (PP=1.0), and revealed a close relationship of isolates from *Sergentomyia minuta* in Europe.

### Median Joining network

To get additional insight into the relationships among the strains, we analyzed our data sets using the Median Joining algorithm network approach. The MJ analysis of three regions was congruent with the topology described in the BI trees, and no haplotype was shared among the species groups.

The network of *cyt b* in Fig 2 (A) showed that the isolate C6 and a *L. donovani* strain from Xinjiang, China (HQ908267) seemed to be the central haplotypes. C1 (MK330198 isolated from two snakes captured at P1 and MK330200 from P2), together with C2 (MK330201 from P3) and C3 (MK330205 from P4) were most closely related to the central HQ908267, within 2 mutational steps. C4 (from P3 and P4) shared the same haplotype with one strain of *L. turanica* from Russia, and only 2 and 3 steps away from the Chinese strains *L. turanica* (from Xinjiang) and *L. gerbilli* (from Gansu) respectively. The *Leishmania sp.* isolate C6 (from P1 and P9) as the central haplotype shared the same sequences to *Leishmania sp.* from China with a wide geographical distribution, including Shandong (HQ908255), Sichuan (HQ908260, HQ908264), Gansu (HQ908263, HQ908271).

For the *Hsp70* region, the haplotype network of Fig 2 (B) could intuitively reflect the genetically small distances between the obtained sequences H1 (MK330208 isolated from P1), H2 (MK330209 and MK330210 from P1 and P2, respectively) and the reference *L. donovani* and *L. infantum* strains from India, Sudan and the northwest of China. The haplotype H5 shared by the isolates from P5 (MK330213) and P6 (MK330214) was the interior haplotype of *Leishmania sp.* and may be older than other tip haplotypes H3 (MK330211) or H4 (MK330212) which were both from P6). It reflected a close distance between JX021439 (Xinjiang, China) and the interior haplotype as three mutational steps.

As shown in the haplotype network of *ITS1*-5.8S in Fig 2 (C), the *Leishmania sp.* group was centered around haplotype KU194967 (Dunhuang, China). All haplotypes found in this group of *Sauroleishmania* isolated from lizards of the northwest of China were very closely related to each other. One strain from Tuokexun (KU194973) shared the same haplotype with I4 (MK300711 from P1). Similarly, two strains from Dunhuang (KU194969, KU194971) and another one from Lukchun (KU194939) shared the same haplotype with I10 (MK300709 from P2). In addition, three strains isolated from *Sergentomyia minuta* of Europe shared a close distance with the *Sauroleishmania* group as less than 3 steps.

## Discussion

This study was the first time to detect the prevalence of *Leishmania* infection in snakes by DNA sequencing. Our result showed that the snakes from 7 loci of the 10 study areas were detected positive for *Leishmania*. Yopurga county, Hami and Dunhuang are the DT-ZVL epidemic foci [5], while Shawan and Texes county are lack of data currently.

Besides one snake (*Natrix tessellate*) from Texes were detected three *Leishmania* species (including *L. donovani*, *L. turanica* and *Leishmania sp.*), the snakes from other positive foci were detected 1-2 species. *Leishmania sp.* was the most widely distributed, while the distribution of the other two species was lack of the significant geographic specificity in this study owing to the limit of the sample size.

The effect of many factors (e.g., the condition of specimen, the DNA extraction methodology, the length of amplified gene sequences and the PCR protocols) would cause different positive rates. Amplification of multiple gene sequences could significantly increase the comprehensiveness of the *Leishmania* infections to be investigated in snakes. Various genetic loci had been developed to detect *Leishmania*, such as kinetoplast DNA (kDNA) genes, nuclear DNA (nDNA) genes and ribosomal RNA (rRNA) genes [25,46,47]. *Cyt b*, one of the cytochromes encoded on the kinetoplast DNA maxicircle, is considered one of the most useful markers that had delimitated most tested *Leishmania* species for phylogenetic work. *Hsp70* gene, an antigen gene on the chromosome, is the most optimal marker for the species discrimination and phylogenetic inference. *ITS1*, one of the highly variable regions of rRNA gene, have been successfully used to resolve taxonomical questions and to determine phylogenetic affinities among closely related Leishmania species. Therefore, three genetic loci were carried out in this study and the positive rates were different indeed. Two samples from Shawan and Tekes were detected *Leishmania sp.* and *L. donovani* by all the three genetic loci while other samples were detected by only one genetic locus. It was suggested that different *Leishmania* primers might have different selection preferences for the gene amplification. At the same time, the use of different primers to amplify the corresponding gene fragments did significantly improve the detection rate of infected snake samples.

It was curious about the spreading mechanism of *Leishmania* in reptiles. Wilson & Southgate assumed that transmission was achieved by the reptile host eating an infected sand fly [48,49]. Nevertheless, there were some reptile-feeding species of *phlebotomine* as we known [50,51]. The same as lizards, it was reasonable that snakes could be infected naturally. Furthermore, there were several reports of snake *trypanosome* [52–54] which was very close to *Leishmania* in phylogeny. In view of the small sample size of each location and species, the infection rate does not make much sense. Nevertheless, all three detected snake species in the endemic foci of VL were found *Leishmania* DNA even the sample size of both *Gloydius halys* and *Natrix tessellate* was 1. Therefore, a wide range of sample size would be necessary for the further study to enrich our understanding of the natural infections of *Leishmania* parasites in snakes.

One of the most noteworthy findings in the present study was that two mammalian *Leishmania* taxa, i.e., *L. donovani complex* and *L. turanica* appeared to occur in snakes. Previously, Zhang et al. had a similar discovery in lizards [19]. They believe lizards played a reservoir role for human leishmaniasis because of the common ancestry of the obtained haplotypes of *L. tropica* and clinical samples. At present, apart from the above study, there are few studies of mammalian parasites natural infections in reptiles. Belova [10] carried out intensive research on reptilian and mammalian *Leishmania* and found that the mammalian parasites (such as *L. donovani*, *L. tropica*, and some other promastigotes recovered from sandflies) were infective to lizards. To determine that the reptile is reservoir host of human leishmaniasis, the *Leishmania* strain collection from VL patients in the epidemic foci and phylogenetic analyses with the reptilian strains are needed.

On the other hand, the *Leishmania sp.* isolated from snakes in this study shared the same haplotypes or located in the close proximity to the strains from VL patients or canines in China as the network shown. Indeed, the same Chinese *Leishmania sp.* isolates were speculated to be closely related to the lizard-infected strains, *L. tarentolae*, on the basis of COII [55] and *cyt b* [28] genes. Analogously, *L. adleri* was found to cause transient CL in humans by Manson-Bahr & Heisch [56], and could infect rodents which the strains were isolated by Adler [57] and Coughlan et al. [58]. *L. tarentolae* was chosen as a candidate vaccine against VL in virtue of the character that the parasites could invade in human macrophages and exist as amastigotes in mammals with slower replication [59]. These studies seem to be contradictory with the former which indicated that *Sauroleishmania* was not pathogenic to human [10]. For the reason that the Chinese strains from patients or canines were isolated in the 70s to 90s of last century, the medical records were no longer available, and we could not confirm whether the patients got leishmaniasis due to *Sauroleishmania* or co-infection with other mammalian *Leishmania*. Thus, to further explore the pathogenicity of *Sauroleishmania*, it is recommended to isolate and culture the promastigotes from liver or blood of reptiles and study the virulence to mammals.

In conclusion, the present study is the first time to investigate the natural Leishmania spp. infection of snakes in the northwest of China. The species identification was primarily achieved by comparisons of the obtained sequences with the GenBank database and determined the belongs with phylogenetic and network analyses, including *Leishmania sp.*, *L. donovani*, *L. turanica*. This is the first time to discover mammalian Leishmania in snakes, after the similar finding in lizards. The *Sauroleishmania* strains in this study closely related to the clinical isolates, which suggests us to be cautious about the widely accepted view that *Sauroleishmania* is nonpathogenic to human. These findings invite us to ponder the importance of snakes in the disease cycle. However, more snake samples in epidemic foci and the pathogenic features of *Sauroleishmania* promastigotes are required for indicating their epidemiological involvement.

## Acknowledgments

We are grateful to Xianguang Guo (Chengdu Institute of Biology, Chinese Academy of Sciences) for his assistance in fieldwork.

